# Effect of dietary zinc supplementation on the gastrointestinal microbiota and host gene expression in the *Shank3B*^−/−^ mouse model of autism spectrum disorder

**DOI:** 10.1101/2021.09.09.459709

**Authors:** Giselle C. Wong, Yewon Jung, Kevin Lee, Chantelle Fourie, Kim M. Handley, Johanna M. Montgomery, Michael W. Taylor

## Abstract

Shank genes are implicated in ~1% of people with autism and mice with Shank3 knock out mutations exhibit autism-like behaviours. Zinc deficiency and gastrointestinal problems can be common among people with autism, and zinc is a key element required for SHANK protein function and gut development. In *Shank3B^−/−^* mice, a supplementary zinc diet reverses autism behaviours. We hypothesise that dietary zinc may alter the gut microbiome, potentially affecting the gut-microbiome-brain axis, which may contribute to changes in autism-like behaviours. To test this, four types of gastrointestinal samples (ileum, caecum, colon, faecal) were collected from wild-type and knock-out *Shank3B^−/−^* mice on either control or supplemented-zinc diets. Cage, genotype and zinc diet each contributed significantly to bacterial community variation (accounting for 12.8%, 3.9% and 2.3% of the variation, respectively). Fungal diversity differed significantly between wild-type and knock-out *Shank3B^−/−^* mice on the control zinc diet, and the fungal biota differed among gut locations. RNA-seq analysis of host (mouse) transcripts revealed differential expression of genes involved in host metabolism that may be regulated by the gut microbiota and genes involved in anti-microbial interactions. By utilising the *Shank3B^−/−^* knock-out mouse model we were able to examine the influence of – and interactions between – dietary zinc and ASD-linked host genotype. These data broaden understanding of the gut microbiome in autism and pave the way towards potential microbial therapeutics for gastrointestinal problems in people with autism.

**Importance:** Previously, supplemental dietary zinc in the Shank3B^−/−^ mouse model of autism spectrum disorder resulted in observations of ASD behaviours reversal; in this study we also used the *Shank3B^−/−^* mouse model to examine the influence of – and interaction between – dietary zinc and ASD-linked host genotype. Sample location along the gastrointestinal tract, genotype and zinc diet explained some of the variation in the microbiota data, with notable bacterial differences between treatment groups. Differential expression of host genes between treatment groups, including antimicrobial interaction genes and gut microbiota-regulated host metabolism genes, suggests that the interplay between gut microbes, the gastrointestinal tract and the brain may play a major role towards the observed amelioration of ASD behaviours seen previously with supplemented dietary zinc. These results widen the scope towards manipulating both dietary zinc and the microbiota itself to ameliorate ASD-related behaviours and associated gastrointestinal issues.

## 1. Background

Autism spectrum disorder (ASD) is a neurodevelopmental condition that includes behavioural impairments in social communication/interaction, as well as stereotypical repetitive behaviours [1]. Recent data indicate that ASD prevalence may be as high as 1 in 54 people [2]. The cause(s) of ASD remain elusive, though it is clear that genetics plays an important role [3, 4], while the potential influence of environmental factors has also come under increased scrutiny [5, 6]. One group of genes that has been implicated in ASD encodes for the <u>SH</u>3 and multiple <u>ank</u>yrin (SHANK) repeat domain proteins. The SHANK protein family represents some of the most important proteins involved in the post-synaptic density scaffold structures in the brain [7, 8], and mutations of the *Shank3* gene are associated with ~1% of all ASD cases [9]. SHANK-related ASD mutations have been generated in animal models of ASD, facilitating their study [10, 11]. For example, the *Shank3B^−/−^* mutant mouse model (ex13-16) displays ASD characteristics, including repetitive behaviours (grooming), anxiety and deficits in social interaction [12].

Gastrointestinal (GI) disorders (ranging from severe constipation to diarrhoea) are common in people with ASD [13, 14]. In at least some cases this may be related to a compromised intestinal epithelial barrier (“leaky gut”) through which dietary products and/or bacterial metabolites may pass to enter into the bloodstream [15–18]. Once in the bloodstream, immune responses and direct interactions with the central nervous system may be elicited. This brings into focus a potential contributing role for the gut microbiota in ASD. The gut-microbiota-brain axis refers to the bidirectional communication between gut microbes and the brain [19–22], and there is evidence from both humans [23, 24] and animal models [25, 26] for a link between the gut microbiota and ASD. For example, Buffington *et al.* (2016; [25]) were able to reverse ASD-associated behaviours and elicit changes in brain neuronal activity in a mouse model of autism by reintroducing the probiotic bacterium *Lactobacillus reuteri*. Numerous studies have reported distinct gut microbiotas (including both bacteria and fungi) in people with and without ASD, but others report no such difference [24, 27–30] and large, well-controlled studies are required to better clarify this relationship. Nonetheless, the malleability of the gut microbiota, through for example diet changes, probiotics or even faecal microbiota transfers between individuals, makes this an attractive option to explore in terms of developing potential microbiota-based therapies for ASD and the associated GI issues.

The *Shank3B^−/−^* knockout (KO) mouse model enables exploration of the interplay between the gut, the gut microbiota and ASD. Indeed, a recent study of the microbiota of *Shank3B^−/−^* mice concluded that there was an imbalance, or dysbiosis, within the gut bacterial community, but that probiotic treatment with *Lactobacillus reuteri* ameliorated ASD-like behaviours [31]. SHANK3 may also play a role in gut permeability changes in ASD, with *Shank3* KO mice exhibiting decreased expression of the tight junction protein ZO-1 in the gut [32].

The micronutrient zinc is required for many processes in the body including GI epithelial barrier function [33], gut permeability [34] and enzyme functionality [35]. Dietary zinc is mainly absorbed in the small intestine, where intestinal mucosa metallothionein proteins, zip proteins and zinc transporters control zinc homeostasis [36–38]. Zinc deficiency has been reported in the ASD population [39–41], potentially due to or leading to compromised nutrient acquisition from an individual’s diet [42], diarrhoea and malnutrition due to reduced integrity of the gut epithelial barrier [43], and increased susceptibility to gut pathogens.

The SAM domain of SHANK3 proteins contains a zinc-binding site, which requires zinc to create structural homomers that scaffold the post-synaptic density [44], and to build post-synaptic protein complexes [45]. Zinc deficiency and loss-of-function *Shank* gene mutations have implications for assembly, maintenance and neurotransmission function at the synapse [46]. Neuronal electrophysiology studies have shown that zinc is responsible for stabilising SHANK3 at the post-synaptic density and regulating excitatory glutamatergic synaptic transmission [45, 47]. Disruption of this zinc-sensitive signalling system has been observed in *Shank3* mutations related to ASD, which may impact the functionality and plasticity of synapses in the brain and potentially result in ASD-like behaviours [45, 48]. Supplementation of zinc in the diet is beneficial in the *Shank3B^−/−^* KO mouse model of ASD, with six weeks of postnatal supplemented dietary zinc resulting in reversal of ASD behaviours as well as changes in the strength and structure of glutamatergic synaptic transmission in the striatum [49]. Moreover, zinc supplementation in pregnant *Shank3B^−/−^* KO mice can prevent the development of ASD-related behaviours in the offspring [8]. Taken together, these studies show the importance of zinc-dependent signalling in not only regulating SHANK at the synapse, but also how this manifests as ASD deficits at the behavioural level.

The link between dietary zinc and ASD behaviour suggests that the gut-microbiota-brain axis is a target for zinc-dependent signalling pathways in ASD. On the basis of this, taken together with multiple indications that gut microbiota influence gut-brain communications, we therefore hypothesised that the synaptic and behavioural changes observed with dietary zinc in *Shank3B^−/−^* KO mice are linked to changes in the gut microbiota, in both bacteria and fungi. The primary aims of this study were to determine the effect of dietary zinc and host genotype on (1) the gut microbiota, and (2) host gene expression in the gut. To evaluate the influence of genotype, we focused on the comparison between wildtype control and *Shank3B^−/−^* KO mice; for dietary zinc, we compared *Shank3B^−/−^* KO mice fed either normal 30 ppm or supplemented 150 ppm dietary zinc. Furthermore, we sought to identify changes in host gene expression within the mouse gut that correlate with dietary zinc treatment.

## 2. Methods

### 2.1 Animals

This study was approved by the University of Auckland’s Animal Ethics Committee (AEC#1299 and AEC#1969) and experimental procedures were in adherence to the ARRIVE guidelines. Shank3^ex13–16−/−^ mice (B6.129-Shank3^tm2Gfng^/J), created by deleting exons 13-16 in the *Shank3* gene [12], were imported from Jackson Laboratories, Bar Harbor, ME, USA and housed in the Vernon Jansen Unit animal facility at the Faculty of Medical & Health Sciences, University of Auckland. Using heterozygous × heterozygous breeding pairs, wild type (WT), heterozygous and homozygous (knockout) offspring mice were created. After weaning (~20 days post-birth), wild-type *Shank3B*^+/+^ mice and mice with a *Shank3B^−/−^* knockout (KO) mutation were randomly assigned to one of two dietary experimental groups: a control zinc diet (30 ppm; [8, 49, 50]) or a supplemented zinc diet (150 ppm), for six weeks (Table 1; Research Diets, Brunswick, NJ, USA (D19410B and D06041101). Four experimental groups were examined: (1) WT30 (wildtype control mice fed 30 ppm zinc); (2) WT150 (wildtype control mice fed 150 ppm zinc); (3) KO30 (*Shank3B^−/−^* mice fed 30 ppm zinc); (4) KO150 (*Shank3B^−/−^* mice fed 150 ppm zinc). Knockout and wildtype mice of the same gender were housed together in individually ventilated cages (to a maximum of 6 mice/cage), with a 12:12 hour light:dark cycle and food available *ad libitum*.

**Table 1.** Faecal and gastrointestinal (gut) sample collection from the four treatment groups.

| Mouse type | Diet |  |
| --- | --- | --- |
|  | Control zinc (30 ppm) | High zinc (150 ppm) |
| Wild type<br>(*Shank3B*<sup>+/+</sup>) | faecal n = 20; gut n = 9 | faecal n = 19; gut n = 10 |
| Homozygous<br>(*Shank3B*<sup>-/-</sup>) | faecal n = 15; gut n = 8 | faecal n = 16; gut n = 10 |

### 2.2 Sample collection and DNA/RNA extraction

After six weeks on the 30 ppm or 150 ppm zinc diet, 9 week old WT and *Shank3B^−/−^* KO mouse faecal samples were collected by placing the mice into a sterile beaker until faecal samples were produced and collected in sterile polypropylene screw-cap tubes and immediately frozen at −20°C. Mice were euthanised by CO_2_, and then gut samples (ileum, caecum, colon) were collected using sterile utensils, and stored in RNA*later* (Ambion, Inc.) in sterile tubes at −20°C.

To extract DNA and RNA, samples were thawed on ice and extracted using the Qiagen AllPrep DNA/RNA Mini Kit (Qiagen, USA), according to the manufacturer’s instructions with minor alterations as follows. Firstly, bead beating was carried out using MP Biomedicals lysing matrix E bead tubes (containing 1.4 mm ceramic spheres, 0.1 mm silica spheres and one 4 mm glass bead) in the Qiagen TissueLyser II at a frequency of 30 Hz for 2 min. All centrifuge steps were adjusted to 11,000 rpm. Extracted RNA was treated with DNaseI (Life Technologies, Auckland, New Zealand) and checked for DNA contamination with PCR (40 cycles) targeting the V3-V4 region of the 16S rRNA gene.

### 2.3 PCR and amplicon sequencing

The hypervariable V3-V4 region of bacterial 16S rRNA genes was amplified using primers 341F and 785R [51] synthesized with an Illumina adaptor for multiplex sequencing (Suppl. Table 1). The fungal Internal Transcribed Spacer 2 (ITS2) genomic region was amplified using the ITS4 and ITS3mix1-5 primers with an Illumina adaptor (Table S1) [52]. Paired-end sequencing generating 2 × 300 bp reads was conducted by Auckland Genomics Ltd (University of Auckland, New Zealand) on an Illumina MiSeq with V3 chemistry.

### 2.4 RNA sequencing

Extracted RNA was sent to Otago Genomics (University of Otago, New Zealand) for ribosomal depletion (Ovation Complete Prokaryotic with mouse AnyDeplete module add-in, Tecan, USA), generation of one equimolar library pool, and paired-end sequencing generating 2 × 125 bp reads, using four lanes of two rapid flow cells on an Illumina HiSeq 2500 with V4 chemistry.

### 2.5 Bioinformatics and statistical analyses

#### 2.5.1 Bacterial amplicon analyses

Amplicon sequence variants (ASVs) were generated from the paired-end sequence files using the DADA2 package v1.14.0 [53] in the R v3.6.1 environment [54]. Sequences were quality filtered, dereplicated and merged, with merged sequences of 400-430 bp retained. Chimeras, singletons and ASVs representing less than 0.00001% of total sequences were removed. The remaining ASVs were taxonomically assigned using the SILVA (v132) database [55]. A rarefaction threshold of 8000 reads/sample was applied, leading to exclusion of two ileum samples and one colon sample from subsequent analyses. R v3.6.1 was used to create statistical graphical output and to analyse data: specifically, by using Permutational multivariate analysis of variance (PERMANOVA) with 999 permutations of the data using the ‘vegan’ package [56] to determine significant sources of variation in microbial data, and by Kruskal-Wallis and Dunn tests [57] with false discovery rate-adjusted p-values to account for multiple pairwise comparisons to identify differentially abundant taxa across treatment groups.

#### 2.5.2 Fungal amplicon analyses

Zero-radius operational taxonomic units (ZOTUs) with 100% sequence similarity (similar to ASVs) were generated using a customised pipeline with USEARCH (v11; [58]). After primer removal, forward reads were trimmed to a maximum length of 220 bp, filtered with a maximum expected error rate of 1 and singletons were removed. The unoise3 algorithm and sintax classifier [59, 60] were used to assign taxonomy with the UNITE reference database (UNITE_22.08.2016; [61]). The ZOTU tables were further refined by removing ZOTUs which did not reach a minimum 0.001% relative abundance threshold across the entire dataset [62]. The data were subsampled to a depth of 500 reads/sample. Analyses were carried out as detailed in section 2.7.1.

#### 2.5.3 RNA sequencing analyses

RNA sequencing files were processed using sickle [63] and trimmomatic [64] to trim adapter sequences and quality trim reads based on a minimum Phred quality score of 30 and a minimum length of 80 bp. Ribosomal RNA reads (ZOTUs silva-bac-16s-id90, silva-bac-23s-id98, silva-arc-16s-id95, silva-arc-23s-id98, silva-euk-18s-id95, silva-euk-28s-id98, rfam-5s-database-id98, rfam-5.8s-database-id98) were removed using SortMeRNA v2.1-gimkl-2017a; [65]). The remaining reads were aligned to the Ensembl 84 mouse reference genome (Mus_musculus.GRCm38,[66]) using the STAR aligner (v 2.7.0e, [67]). Reads were then quantified using the feature counts software [68] and analysed using R (v3.6.1). The R package edgeR [69] was used to identify differentially expressed genes, with relationships among samples visualised using multi-dimensional scaling (MDS) plots generated in R. Specific gene analysis was carried out using transcripts per million (TPM) and the KEGG database [70].

### 2.6 Accession numbers

The sequence data from this study have been deposited in the NCBI Sequence Read Archive under accession number SUB9937488.

## 3. Results

### 3.1 Influence of mouse genotype and dietary zinc supplementation on bacterial and fungal communities

For bacterial analyses, 70 faecal and 111 gastrointestinal samples (representing 37 ileum, caecum and colon samples per mouse) were sequenced, resulting in 24,619,426 16S rRNA gene sequence reads. After rarefying to 8,000 reads/sample, 178 (70 faecal, 35 ileum, 37 caecum, 36 colon) samples were retained and we observed 375 ASVs classified into seven bacterial phyla and 45 genera. For fungi, we sequenced 139 (33 ileum, 33 colon, 33 caecum and 40 faecal) samples, resulting in 3,909,164 ITS2 reads. After rarefying to 500 reads/sample, 57 (13 ileum, 12 caecum, 17 colon and 15 faecal) samples were retained and we observed 65 ZOTUs classified into three phyla and 11 genera.

Overall, there was considerable overlap in bacterial (Figure 1) and fungal (Suppl. Figure 1) community compositions among the four experimental groups (WT mice fed 30 ppm zinc, WT fed 150 ppm zinc, KO *Shank3B^−/−^* mice fed 30 ppm, KO *Shank3^−/−^* mice fed 150 ppm). Of bacterial 16S rRNA gene sequences that could be assigned to phylum level, the dominant phyla across the entire dataset were *Verrucomicrobia* (35.3 mean ± 7.8% S.D.), *Bacteroidetes* (31.9 ± 6.2%) and *Firmicutes* (22.2 ± 8.4%) (Figure 1A). The genus *Akkermansia* dominated across the entire dataset regardless of genotype or dietary zinc level (35.0 mean ± 8.1% S.D.), with *Faecalibaculum* (6.8 ± 6.5%), *Muribaculum* (4.4 ± 3.6%), *Escherichia*/*Shigella* (4.3 ± 2.6%) and *Dubosiella* (4.1 ± 3.8%) also abundant (Figure 1B). Although the majority of the detected fungal ZOTUs (67.3 mean per sample ± 39.9% S.D.) could not be classified beyond kingdom level, the phylum *Ascomycota* was dominant among those taxa that could be assigned (20.1 ± 31.9%), with other ZOTUs classified to the phyla *Basidiomycota* (9.6 ± 25.7%) and *Zygomycota* (3.0 ± 8.1%) (Figure S1). At genus level, the fungi *Rhodotorula* (6.4 ± 22.4%) and *Cyberlindnera* (6.2 ± 20.0%) were most prevalent among those identified, with nine other genera ranging from 0.01-2.70% relative abundance across the data set (Suppl. Figure 1B). The lack of distinct microbial signatures between the four experimental groups, as shown by the taxonomic bar charts, was also evident in the non-metric multidimensional scaling plots for both bacteria and fungi (Figures 2A and 2B). Nevertheless, there was marked similarity among gut section communities, with the fungal faecal communities specifically notably differing along the gastrointestinal tract (Figure 2B).

**Figure 1.**
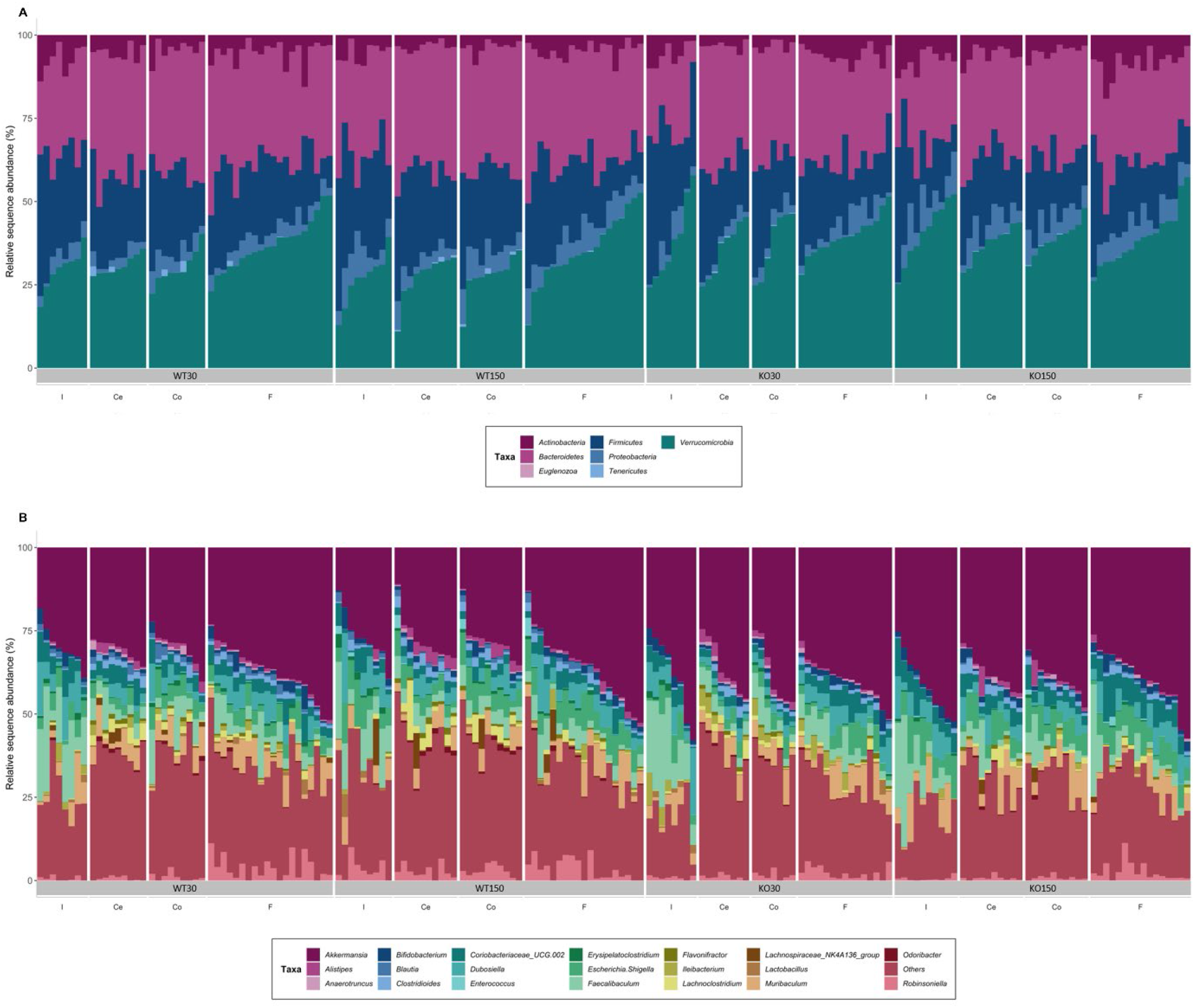
16S rRNA gene-based taxonomic summary plot of the relative abundance of bacterial phyla (A) and top 20 most abundant genera (B) in each of the four treatment groups. (Wild type control zinc diet WT30 n = 46, wild type supplementary zinc diet WT150 n=48, Shank3B^−/−^ KO control zinc diet KO30 n= 38 and Shank3B^−/−^ KO supplementary zinc diet KO150 n=46). All other genera and ‘unknown’ genera taxonomic assignments are grouped in ‘Others’. Gastrointestinal sections are noted as I (ileum), Ce (caecum), Co (colon), F (faecal).

**Figure 2.**
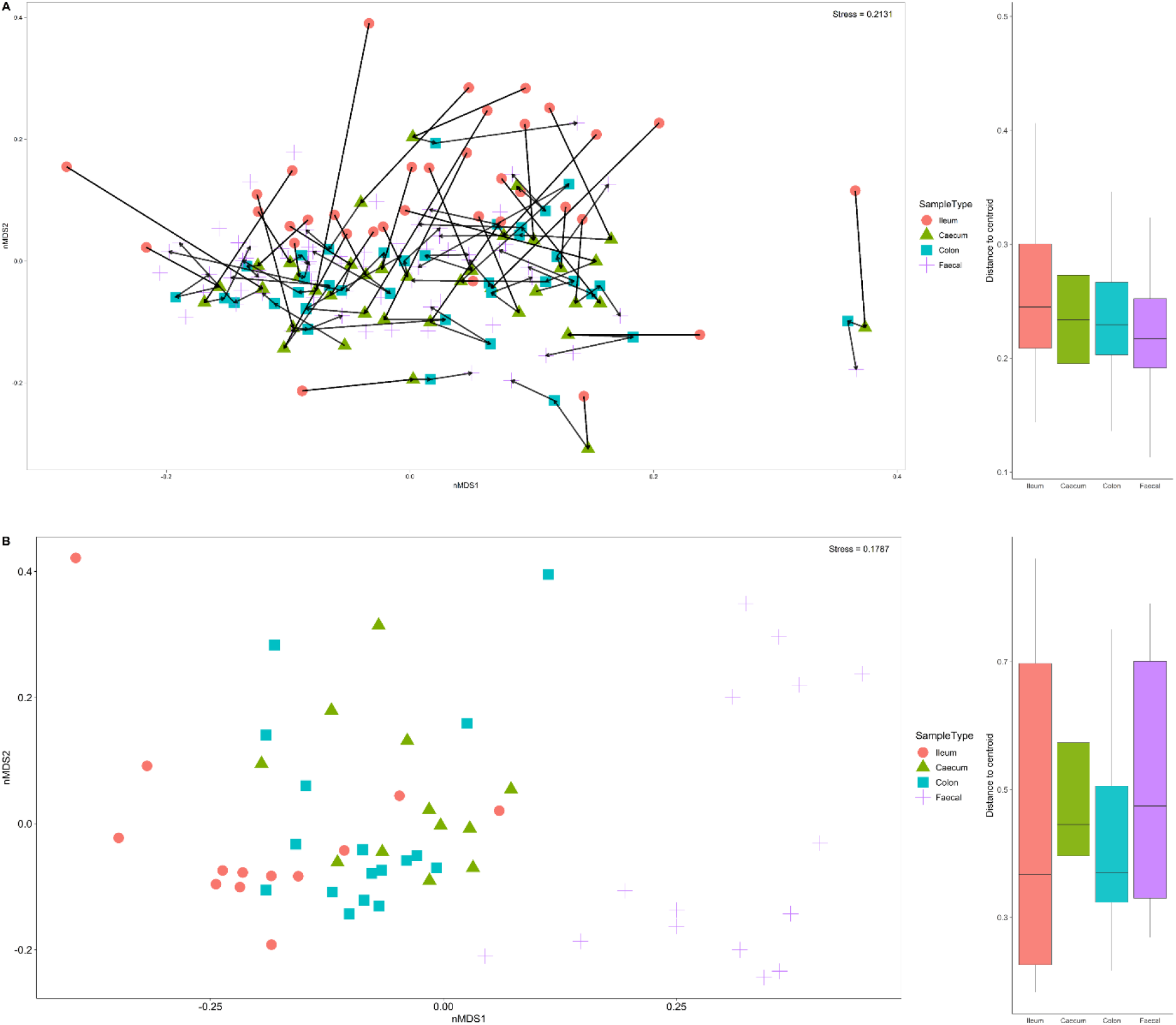
Beta-diversity Bray-Curtis dissimilarity non-metric multidimensional scaling plot from all samples with indicated sample type (ileum, caecum, colon and faecal) based on bacterial (A) and fungal (B) communities. The arrow in (A) depicts the sequential order through the gastrointestinal tract for an individual mouse. The microbial community within a sample is represented as a point: the closer together the points the more similar, the further apart the more dissimilar. Box and whiskers plots accompany the nMDS plots showing the distance from the centroid box and whisker plots of different sample types.

Bacterial community richness and diversity varied among samples, with 74 to 148 distinct bacterial ASVs observed per sample (Figure 3A). The caecum samples (based on Jost1, Shannon and Inverse Simpson) and faecal samples (based on richness) had the highest alpha-diversity, depending on the diversity measure applied. When aggregating samples from all gut locations for a given experimental group, there was significantly higher (p<0.05) bacterial diversity in the WT compared with the *Shank3B^−/−^* KO mice (Figure 3A), irrespective of alpha-diversity metric and zinc diet. For the fungal ITS2 data, significantly higher diversity – supported by all four alpha-diversity metrics – was observed between the WT30 group (control mice fed 30 ppm zinc), compared with KO30 (*Shank3B^−/−^* KO mice also fed 30 ppm zinc) group (Figure 3B), showing that gut fungal diversity is significantly affected in *Shank3B*^−/−^ KO mice. In contrast, there were no significant differences between the *Shank3B*^−/−^ KO mice fed the supplemented zinc diet (KO150) and the WT30 controls, suggesting that dietary zinc may revert fungal diversity to that observed in WT mice.

**Figure 3.**
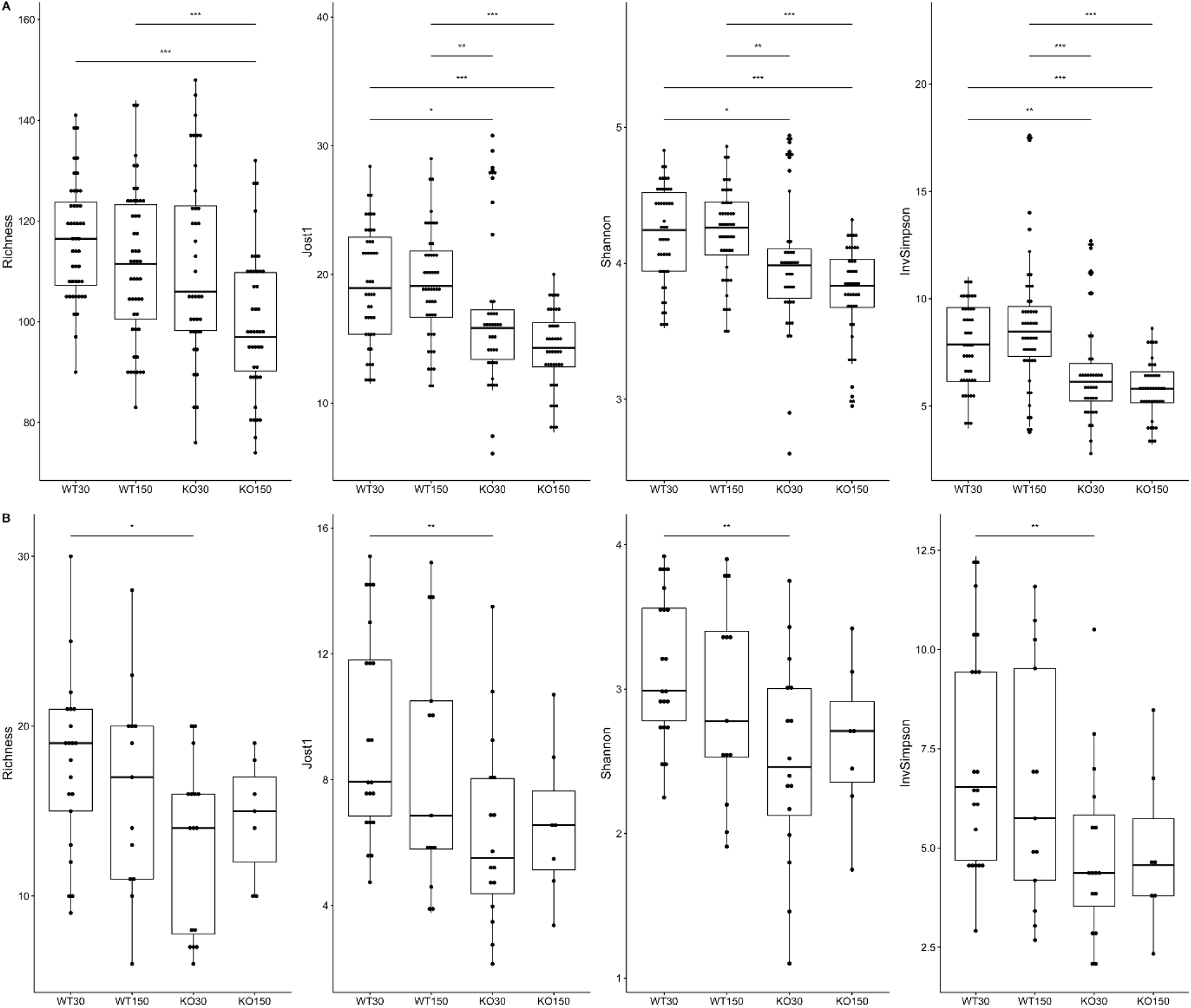
Alpha diversity box and whisker plots based on Richness (observed taxa), Jost1, Shannon and Inverse Simpson index for bacteria (A) with WT30 n = 46, WT150 n =47, KO30 n = 38, KO150 n = 43 and fungi (B) WT30 n = 21, WT150 n =13, KO30 n = 16, KO150 n =7. * p ≤ 0.05, ** p ≤ 0.01, ***p ≤ 0.001.

To identify potential factors contributing to the observed statistically significant differences in beta diversity, we performed PERMANOVA analyses to determine whether caging, genotype, dietary zinc level, or gut sample location also play a role. For bacteria, individual cage contributed the greatest amount to variation within the microbiota data, accounting for 12.8% (*p*=0.001) of observed variation (Table 2). Gut sample location, mouse genotype, dietary zinc level and the genotype:zinc diet interaction were also significant (*p*≤0.008), explaining 11.4%, 3.9%, 2.3% and 1.1% of microbiota variation, respectively.

**Table 2.** PERMANOVA of bacterial ASV data table modelling cage number and the interaction between genotype and zinc diet, 999 permutations. Sample type refers to sample location along the gastrointestinal tract.

|  | Df | SumsOfSqs | MeanSqs | F.Model | R<sup>2</sup> | Pr(>F) |
| --- | --- | --- | --- | --- | --- | --- |
| Cage | 1 | 1.5843 | 1.5844 | 31.8620 | 0.1284 | 0.0010 |
| Sample type | 3 | 1.4028 | 0.4676 | 9.4040 | 0.1137 | 0.0010 |
| Genotype | 1 | 0.4803 | 0.4803 | 9.6580 | 0.0389 | 0.0010 |
| Zinc diet | 1 | 0.2883 | 0.2883 | 5.7970 | 0.0234 | 0.0010 |
| Genotype:Zinc diet | 1 | 0.1324 | 0.1324 | 2.6630 | 0.0107 | 0.0080 |
| Residuals | 170 | 8.4533 | 0.0497 | 0.6850 |  |  |
| Total | 177 | 12.3414 | 1.0000 |  |  |  |

Bray-Curtis dissimilarity non-metric ordination plots showed that bacterial communities associated with ileum samples were overall less similar compared to the caecum, colon and faecal samples within a mouse individual. Despite the high overall similarity of bacterial faecal communities to caecum and colon communities in the dataset as a whole, when considering each individual mouse, caecum and colon communities were the most similar (Figure 2A). A broadly similar trend was observed for the fungal data, whereby dietary zinc levels explained 3.7% of observed variation overall (*p*=0.007), while gut sample location accounted for almost one-quarter (23.4%, *p*=0.001) of variation (Table 3; Figure 2B); these results reflect substantially greater discrimination of faecal fungal communities based on gut location than observed for bacteria.

**Table 3.** PERMANOVA of fungal ZOTU data table modelling zinc diet and sample type, with 999 permutations. Sample type refers to sample location along the gastrointestinal tract.

|  | Df | SumsOfSqs | MeanSqs | F.Model | R<sup>2</sup> | Pr(>F) |
| --- | --- | --- | --- | --- | --- | --- |
| Zinc diet | 1 | 0.6988 | 0.6988 | 2.6457 | 0.03709 | 0.007 |
| Sample type | 3 | 4.4062 | 1.4687 | 5.5609 | 0.23389 | 0.001 |
| Residuals | 52 | 13.7340 | 0.2641 |  | 0.72902 |  |
| Total | 56 | 18.8390 |  |  | 1.0000 |  |

### 3.2 Specific microbial taxa as biomarkers for experimental groups

We identified specific microbial taxa that were differentially abundant among the four experimental groups, using Kruskal-Wallis rank-sum and Dunn testing with adjustments for multiple pairwise comparisons. A total of 74 bacterial ASVs differed significantly in abundance between the WT30 controls and the KO30 groups (Table S2), with the majority classified as members of the phyla *Bacteroidetes* and *Firmicutes*, and the families *Muribaculaceae* and *Lachnospiraceae*. Individual ASVs with the lowest *p*-values (*p*<3.78 × 10^−9^) were ASVs 182/155/135 (all *Akkermansia*), ASV 90 (*Lachnoclostridium*) and ASV 69 (*Muribaculaceae*). Sixty-five ASVs differed significantly (*p*<0.05) in relative abundance between the KO30 and KO150 groups, with 65 ASVs also differing between the WT30 and KO150 groups, including the bacterial taxa described earlier in this paragraph (Table S2). Across all experimental groups, 24 fungal ZOTUs were identified as being differentially abundant (Table S3). Only five of these ZOTUs could be assigned to below kingdom level: *Rhizopus* ZOTUs 5, 7 and 502, and *Aspergillus* ZOTUs 11 and 19 (Table S3).

### 3.3 Effect of mouse genotype and dietary zinc supplementation on host gene expression

In total, the transcriptomes of 24 (12 ileum, 12 colon) samples were sequenced, resulting in 787,500,000 reads. Following quality control, trimming and removal of ribosomal RNA, 787,447,474 reads remained. For each sample, 60-76% (mean = 7,394,185) of reads mapped to the mouse genome. A single colon sample failed QC and was removed from subsequent analyses.

Gene expression patterns correlated strongly with sample location. Compared to the colon, ileal samples exhibited lower expression of functional gene groups related to KEGG categories Digestive system, Cell motility and Metabolism of other amino acids, while genes relating to energy metabolism were typically more highly expressed in the colon (Figure 4). KEGG functional groups of genes related to Signal transduction, Endocrine and metabolic diseases, and Neurodegenerative diseases were particularly highly expressed in all colon and ileum samples, encompassing ~ 86 - 89% of all transcripts (KEGG Level 2, Figure 4). Overall, there were KEGG group expression differences between the WT and *Shank3B*^−/−^ KO genotypes as well as between the 30 ppm and 150 ppm zinc diets within a genotype, with differences most notable among colon samples. Genes involved in energy metabolism were enriched in the colon of *Shank3B*^−/−^ KO mice (on both zinc diets) compared to wild type mice on the control zinc diet. Additionally, immune disease genes had decreased expression on the supplemented zinc diet (150 ppm) compared to the control zinc diet (30 ppm) in the colon (Figure 4).

**Figure 4.**
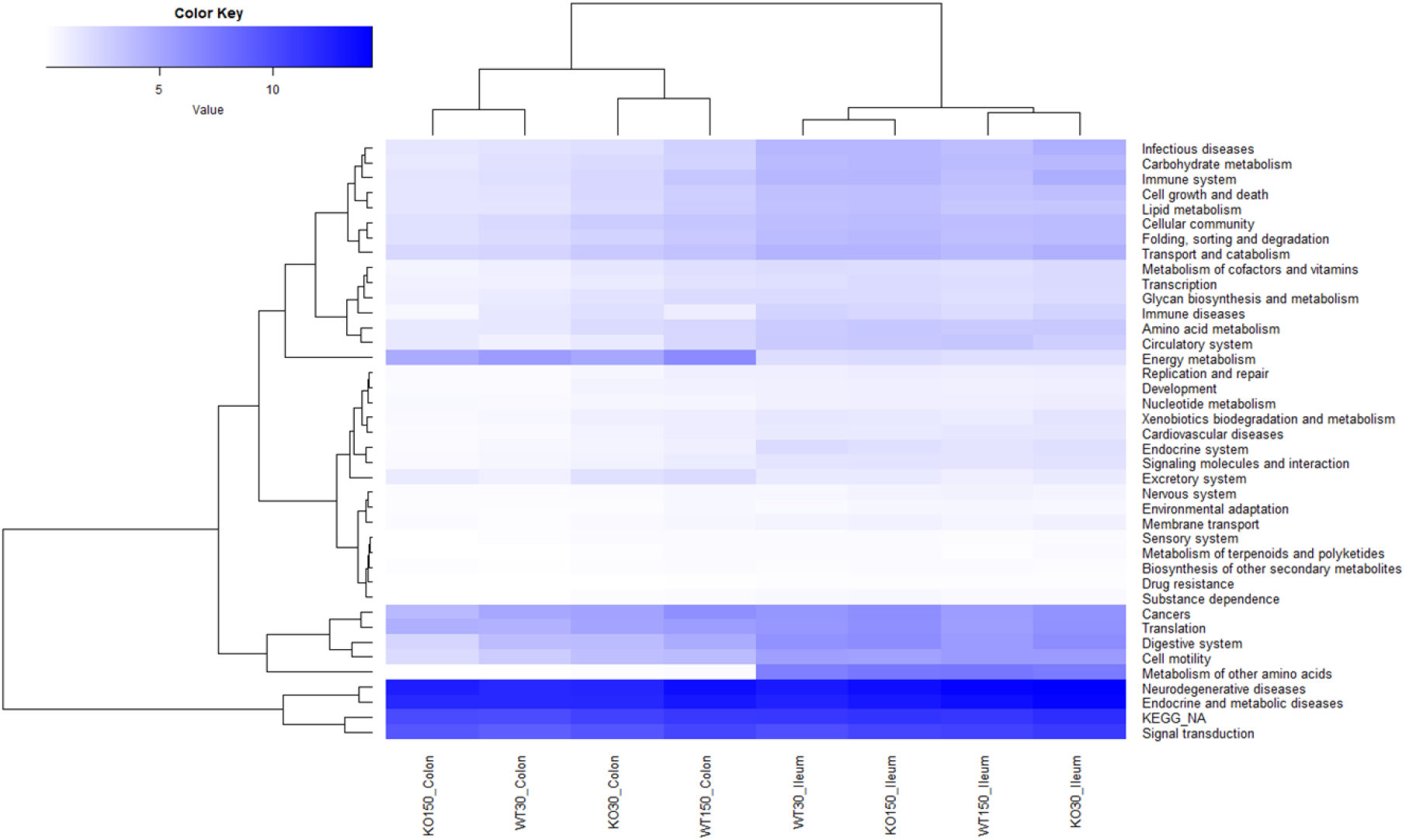
Heatmap of KEGG orthologous groups (level 2) expressed by the host (mouse) across the four treatment groups and two sample types (ileum and colon). RNA-seq data were normalized to total counts and presented as log +1 transcripts per million. Samples within a treatment group and sample type were pooled together for analysis as biological replicates.

Across the entire data set, a total of 1,387 genes were significantly differentially expressed based on differential expression analysis. Consistent with the clustering shown in Figure 4, ileum and colon samples were separated in two-dimensional space in MDS plots (Figure 5A), with greater diversity in the expression of these genes in the colon. By contrast, distribution of samples in the ordination did not correlate with experimental group, which encompasses zinc diet and mouse genotype (Figure 5B). When differential expression analyses were carried out on each of the two gut sample locations (with data from the four experimental groups combined per location), 1,271 genes among the ileum samples and 741 genes among the colon samples were significantly differentially expressed (data not shown). Many of these are related to antimicrobial interactions (alpha-defensins, mucin 2, carbonic anhydrase 1, intelectin) and host metabolism genes that may be regulated by gut microbiota (e.g. NADH dehydrogenases).

**Figure 5.**
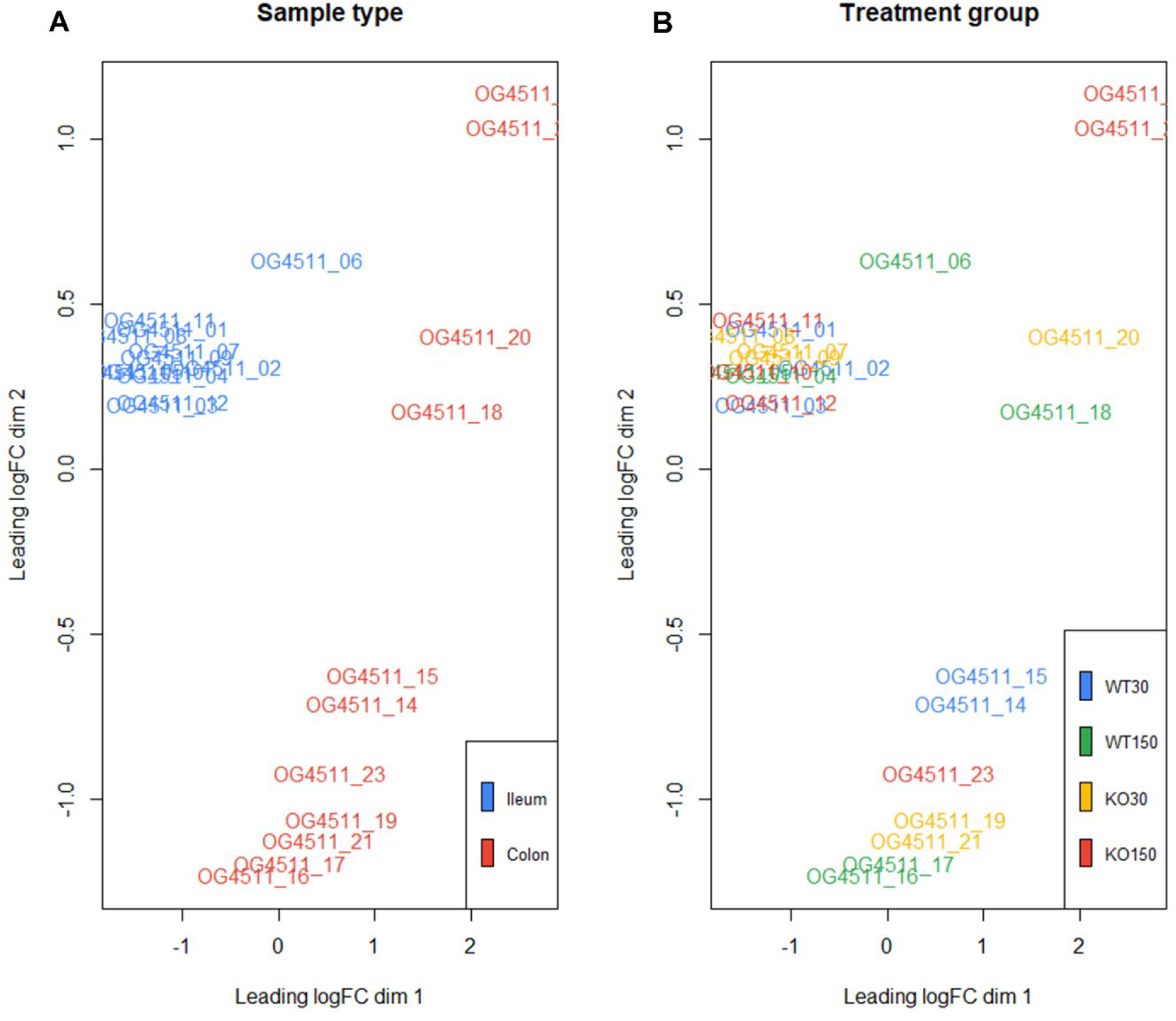
Multi-dimensional scaling plots of the samples with distances showing leading logFC (the average of the largest absolute log fold changes between each pair of samples) coloured by sample type (A) and treatment group (B).

To investigate non-parametric differential feature (gene expression) abundance between treatments, linear discriminant analyses of effect size (LEfSe) was used. This identified 54 genes (biomarkers) that were significantly differentially expressed between the KO30 and KO150 groups (Figure 6A). For this analysis, ileum and colon samples for each experimental group were combined to find key differences regardless of sample type. NADH dehydrogenases, DNA/RNA binding proteins, zinc finger and transmembrane proteins were among the most differentially expressed gene categories. We focused our subsequent analysis on two main categories: alpha defensins, due to their importance in the control of bacteria at the mucosal barrier, and genes related to tight junction proteins, due to their integral role in mediating gastrointestinal permeability [71]. Overall, alpha defensin genes were more highly expressed in the ilea of *Shank3B^−/−^* (KO) mice - with slightly increased expression in the supplemented zinc diet KO mice - compared to the ilea of wild-type mice (Figure 6B). Expression of tight junction genes was significantly lower in the colon samples from *Shank3B^−/−^* KO mice receiving zinc supplementation compared to their wild-type counterparts on the same diet (*p =*0.0139), while the ileum samples showed the opposite trend with greater expression in *Shank3B^−/−^* KO mice than wild type mice (Figure 6C).

**Figure 6.**
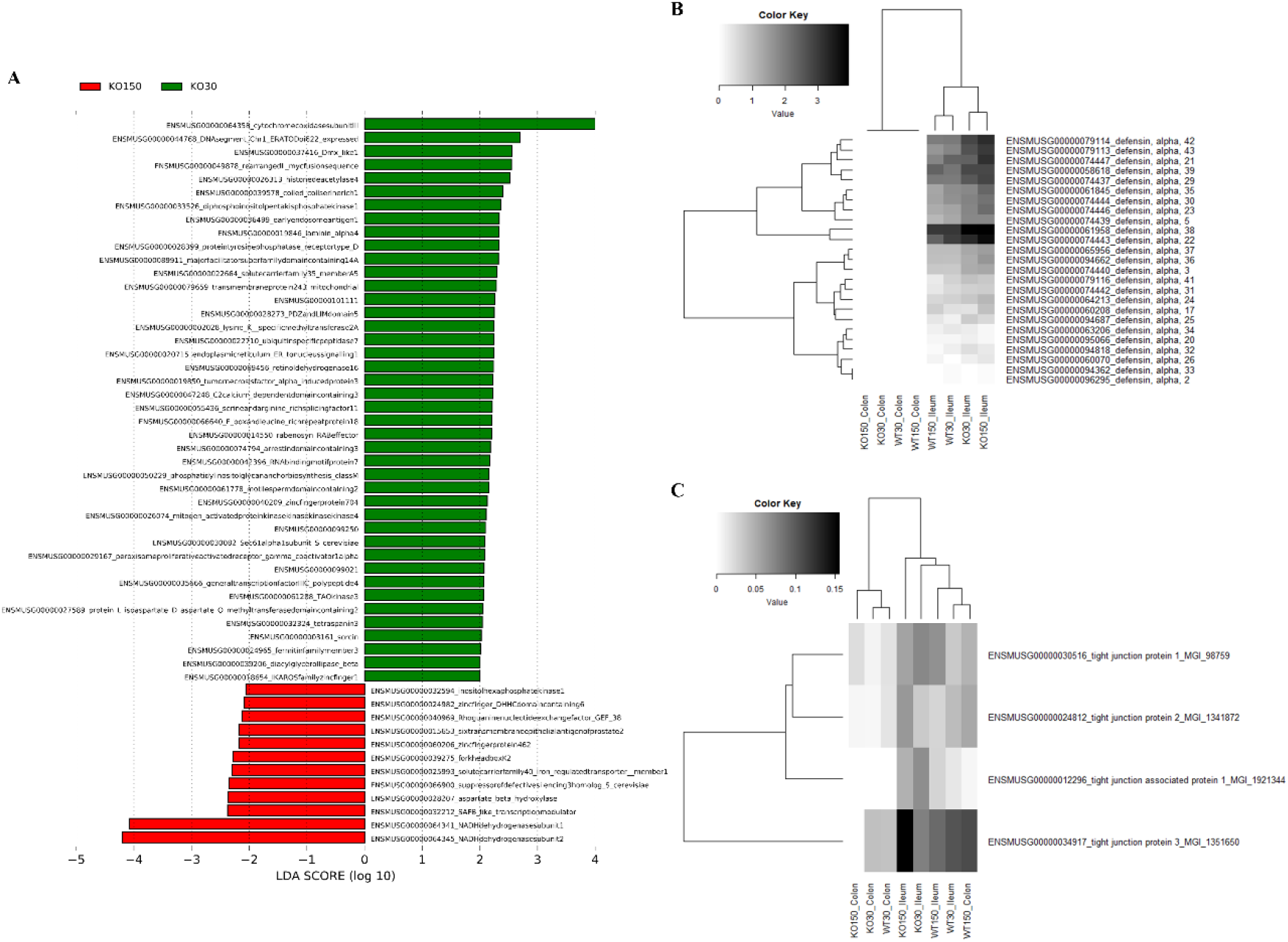
RNA-seq analyses of host (mouse) gene expression, LEfSe analyses identifying biomarker differences between KO30 vs KO150 for both ileal and colon samples combined (A). The higher the LDA score the larger the effect size, with all genes presented differing significantly between the treatment groups. Heatmap of alpha defensin genes (B) and tight junction protein genes (C) across the four treatment groups and two sample types. RNA-seq data were normalized to total counts and presented as log +1 transcripts per million. Samples within a treatment group and sample type were pooled together for analysis as biological replicates.

## 4. Discussion

The role of the gut microbiota in ASD has come under considerable scrutiny in the past decade, with evidence for its importance coming from studies of both human populations and animal models [24–26]. Neural, immunological and hormonal pathways all offer potential mechanisms by which gut microbes may influence neurological development. Changes in gut microbial composition and resulting functional differences have been observed with differential dietary zinc intake in porcine and murine models as well as human studies [72–74]. We recently described dietary zinc-induced reversal of ASD behaviours in *Shank3B^−/−^* KO mice [8, 49]. Zinc plays a critical role in gut health and, taken together with published data linking the gut-microbiota-brain axis to ASD, these observations raise the intriguing possibility that the effects of dietary zinc on ASD could be linked to changes in the gut microbiota. In this study, we therefore examined the effects of dietary zinc and host genotype on (1) the gut microbiota in *Shank3B^−/−^* KO mice, and (2) host gene expression in the gut.

### 4.1 Zinc supplementation and host genotype influence microbiota composition

When considering the gut-associated microbiota, differences among individual mice were substantial. Even within a given dietary zinc level or genotype group, the phylum- and genus-level bacterial taxonomic compositions varied greatly, albeit with overall dominance by certain taxa, most notably the genus *Akkermansia* (phylum *Verrucomicrobia*). Despite such inter-individual variation, which is consistent with previous autism gut microbiota studies [75], our data show significant effects of host genotype, dietary zinc level, and their interaction. Specifically with genotype for both bacterial and fungal data, across all alpha-diversity indices wild type mice displayed significantly increased diversity compare with *Shank3B^−/−^* KO mice.

The clearest differences related to dietary zinc levels and/or genotype were evident not at the level of entire bacterial communities, but rather at the level of specific ASVs, consistent with other gut microbiome studies [76, 77]. For example, 74 ASVs, mostly from the families *Muribaculaceae* (*Bacteroidetes*) and *Lachnospiraceae* (*Firmicutes*), were significantly differentially represented between the WT and KO groups that were both fed 30 ppm zinc. When comparing the two genotypes on the control zinc diet, *Muribaculaceae*, *Blautia* and *Erysipelotrichaceae* were more abundant in WT30 mice than KO30 mice. *Blautia* produces butyrate, a short-chain fatty acid (SCFA), and is beneficial for host health. Butyrate has positive anti-inflammatory effects at the mucosa and fortifies the epithelial barrier, with implications for the gut-microbiota-brain axis as it is able to cross the blood-brain barrier and produce neuromodulatory effects [78–80]. Other SCFAs such as acetate, propionate and succinate can also be produced by members of the family *Muribaculaceae*. This family represents one of the largest proportions of gut bacteria found in mice, and has been suggested to play a similar role to the genus *Bacteroides* in humans, occupying similar niches and metabolising the same dietary source - complex carbohydrates [81, 82]. The balance of host and gut microbiome requirements for zinc is essential for both symbiotic systems to survive. Microbial high-affinity membrane transport systems and zinc binding proteins for competitive uptake of zinc between gut microbes are crucial for their persistence and growth along the gastrointestinal tract.

Among the *Shank3B^−/−^* KO mice, the bacterial families *Enterobacteriaceae* and *Coriobacteriaceae* were the top two significant biomarkers that were more abundant in the supplemented zinc diet (KO150) mice compared with the control zinc diet (KO30) mice. KO30 mice, instead, had increased abundances of *Muribaculaceae* and *Ileibacterium*. Diet changes can alter the host production of bile acids, which can shift the gut community in favour of organisms that can metabolise bile acids such as members of the *Coriobacteriaceae* [83]. This, in turn, can affect gut barrier permeability by interaction of bile acids with the epithelial cells [84]. Similarly, members of *Muribaculaceae*, *Blautia* and *Faecalibaculum* were more abundant in WT30 mice than KO150, which again had higher abundances of *Coriobacteriaceae* and *Escherichia/Shigella*.

It is worth considering the extent to when the gut microbiota may be directly utilising zinc from the diet, potentially in competition with the host animal. Smith *et al.* (1972; [85]) estimated that gastrointestinal bacteria utilise 20% of the dietary zinc consumed in a germ-free/pathogen-free rat study [85]. By zinc supplementation in the host diet (150 ppm), this potentially results in increased zinc availability along the gastrointestinal tract for the microbial community, affecting its overall community structure and function.

While our main focus was on the influence of dietary zinc on the gut microbiota of *Shank3B^−/−^* KO mice, this study also provides an opportunity to revisit the association between host genotype and microbiota composition in this animal model. An earlier study by Tabouy and colleagues described reduced bacterial alpha-diversity in *Shank3B^−/−^* KO compared with wild-type mice, as well as differences in the relative abundances of several bacterial genera including *Lactobacillus* and *Prevotella* [31]. The authors interpreted these differences as reflective of a dysbiosis, or dysregulation, of the microbiome. We also observed differences of a comparable magnitude between the *Shank3B^−/−^* KO and wild-type mice microbiotas (on the normal 30 ppm zinc diet), specifically a decrease in bacterial diversity in KO mice, but contend that these do not necessarily constitute a dysbiosis. Indeed, there were marked similarities at both phylum and genus levels between the *Shank3B^−/−^* KO and wild-type bacterial communities in our data (Figure 1), as well as in the aggregated genus-level data of Tabouy and colleagues (Supplementary Figure 2 in their paper). Moreover, while statistically significant, alpha-diversity in the female mice (which displayed the strongest effect) still only decreased by <10% in the Tabouy *et al*. data set. Widespread use of the term “dysbiosis” to describe any shift in microbial communities has been criticised [86] and we suggest that this term would more appropriately apply to a situation of extreme microbiota depletion, such as seen in *Clostridioides difficile* infection [87], rather than the more subtle changes evident in both our data and those of Tabouy *et al*. [31]. Nonetheless, whether or not the microbiota differences observed here and by Tabouy et al. [31] constitute a dysbiosis *per se*, the functional implications of reduced gut-associated microbiota diversity in *Shank3B^−/−^* KO mice warrant further investigation.

It is important to note that genotype and dietary zinc factors explained ~7% of variation in the bacterial biota, compared with almost 13% attributable to individual cage differences. The explanatory power of dietary zinc and genotype was therefore lower, despite striking host behavioural differences associated with zinc supplementation [49], which were not influenced by mouse cage assignment.

Our data also show significant effects of genotype and dietary zinc on fungal diversity. Twenty-four fungal ZOTUs differed significantly in relative abundance across the entire dataset. The five that could be assigned to below kingdom level represented the genera *Rhizopus* and *Aspergillus.* We observed that, on the control diet, KO mice displayed significantly lower fungal diversity compared with WT mice, and moreover that dietary zinc treatment in KO mice reverted fungal diversity towards WT levels. Zinc is key for fungi and has been demonstrated to enhance growth of *Rhizopus* with higher zinc levels [88] and *Aspergillus* [89] by stimulating RNA synthesis and transcription. Recently a child with ASD treated with antifungals to target the removal of *Rhizopus* reversed their autism behaviours, though the efficacy of this treatment for a wider population is yet to be tested [90]. Mycotoxins and fungi have been postulated to be involved in the pathobiology of ASD due to their high prevalence in the urine and blood of children on the spectrum, but this is largely understudied [91, 92]. Potential targets for antifungal treatment include zinc transporters (due to their ubiquity and importance in fungi) as well as probiotic treatments such as *Lactobacillus* aimed at adsorbing mycotoxins or changing gut microbial communities [93–95]. Therefore, dietary zinc supplementation may aid in maintaining the balance between a healthy fungal community and their interactions with the *Shank3B* deficient host which ultimately may affect their phenotype. Together with our data, these findings highlight fungal diversity and zinc as an important focus for future studies.

### 4.2 Dietary zinc supplementation has a minor but measurable effect on host gene expression in the mouse gut

Consideration of host-microbe interactions demands attention to not only the microbiota, but also the host itself. We therefore utilised RNA-Seq to investigate patterns of host (mouse) gene expression in the ileum and colon of both WT and *Shank3B^−/−^* KO mice fed either 30 ppm or 150 ppm dietary zinc. As overall gene expression correlated most strongly with gut region (ileum *vs* colon) rather than experimental group (dietary zinc or genotype) (Figure 5), we focused our attention on those genes that were significantly differentially expressed and/or directly relevant to integrity of the gut epithelial barrier.

Alpha defensins are antimicrobial peptides active against a wide range of microorganisms [96]. Consistent with their known prevalence in the small intestine [97, 98], we documented higher expression of alpha defensin genes in the ileum compared with colon samples. Expression of several alpha defensins was higher in the *Shank3B^−/−^* KO mice compared to the wild-type mice, especially those on the supplemented zinc diet. Whilst speculative, this could indicate enhanced antimicrobial activity in *Shank3B^−/−^* KO mice, and that this is further enhanced by high dietary zinc. This suggests that the gut microbiome may be more strictly controlled in *Shank3B^−/−^* KO mice compared to wild type mice by various host-microbe interactions.

Given the significant research focus on potential leaky gut in ASD, we also focussed our analysis on the expression of tight junction proteins between WT and *Shank3B^−/−^* KO mice fed normal or supplemented zinc levels. Tight junction proteins are crucial for integrity of the gut epithelium, preventing potentially harmful microorganisms or their metabolites from leaking out of the gut and into the bloodstream. Zinc supplementation enhances and maintains the intestinal barrier by regulating expression of several tight junction proteins [99]. Tight junction expression in the ileum samples from *Shank3B^−/−^* KO mice was higher than in the corresponding WT30 samples, which may suggest the presence of intestinal inflammation or compromised barrier integrity that can ultimately impact the gut-microbiota-brain axis [100]. Dietary zinc supplementation correlated with higher expression, regardless of host genotype, particularly in the ileum. This is intuitive given the greater absorbance of dietary zinc in the small intestine [36–38]. Overall, our observations suggest that zinc may be playing a role in the reinforcement of the gut epithelial barrier that may help to tightly regulate the gut-microbiota-brain axis and perhaps ameliorate the resulting ASD behaviours.

### 4.3 Gut region correlates more strongly with fungal than bacterial community composition

We sampled the mouse gut at several regions along the gastrointestinal tract, namely the ileum, caecum and colon, as well as faecal samples. For bacteria, there was no clear separation of microbiota composition related to gut region in the ordination plot (Figure 3A), although for many individual mice the ileum samples were somewhat distinct. There was much greater separation for fungi, with the fungal composition of faecal samples being markedly different from that of other sample types (Figure 2B) and greater variation explained in the PERMANOVA models (23.39%, *p*=0.001) compared to the bacterial data (11.37%, *p*=0.001). The bacterial community was substantially more diverse than fungal communities across the dataset (Figure 3), therefore elucidating bacterial and fungal functions and interactions as a community is key to determine their interplay between the host and gut microbial community.

## 5. Conclusions

To our knowledge this study is the first to investigate the effect of dietary zinc supplementation on the gut microbiota in the context of autism. By utilising the *Shank3B^−/−^* KO mouse model we were able to examine the influence of – and interaction between – dietary zinc and ASD-linked host genotype. We found that sample location along the gastrointestinal tract, genotype and zinc diet explained some of the variation seen in the microbiota data, with notable bacterial ASV differences between treatment groups. While we observed statistically significant differences between the alpha-diversity of microbiotas of different genotype and dietary zinc groups, we argue that these do not necessarily constitute a major breakdown, or dysbiosis, of the gut microbiota. Differential expression of host genes among the treatment groups, including antimicrobial interaction genes as well as gut microbiota-regulated host metabolism genes, suggests that the interplay between gut microbes, the gastrointestinal tract and the brain may play a major role towards the observed amelioration of ASD behaviours seen previously with supplemented dietary zinc. Ongoing research in our group will address effects on the functional potential of these microbial communities as well as the scope for manipulating both dietary zinc and the microbiota itself to ameliorate ASD-related behaviours and associated gastrointestinal issues.

## Ethical Approval and Consent to participate

This study was approved by the University of Auckland’s Animal Ethics Committee (AEC#1299 and AEC#1969) and experimental procedures were in adherence to the ARRIVE guidelines.

## Availability of data and materials

The datasets supporting the conclusions of this article are available in the NCBI Sequence Read Archive repository under accession number SUB9937488 https://www.ncbi.nlm.nih.gov/sra/PRJNA747052.

## Authors’ contributions

JMM provided the design of this study. GCW collected samples and analysed all the data. GCW (main) and MWT wrote the manuscript. YJ, KL and CF undertook mice husbandry/wellbeing. KMH provided advice on methodology and analysis. GCW, MWT, KMH, JMM all edited the manuscript. All authors have read and approved the final manuscript.

## Acknowledgements

The authors would like to thank the Auckland Genomics Centre, The University of Auckland, Auckland, New Zealand for assistance with microbial amplicon sequencing and the Otago Genomics Facility, University of Otago (Dunedin, New Zealand) for facilitating the RNA sequencing.

## Supplementary Material

**Table S1.**
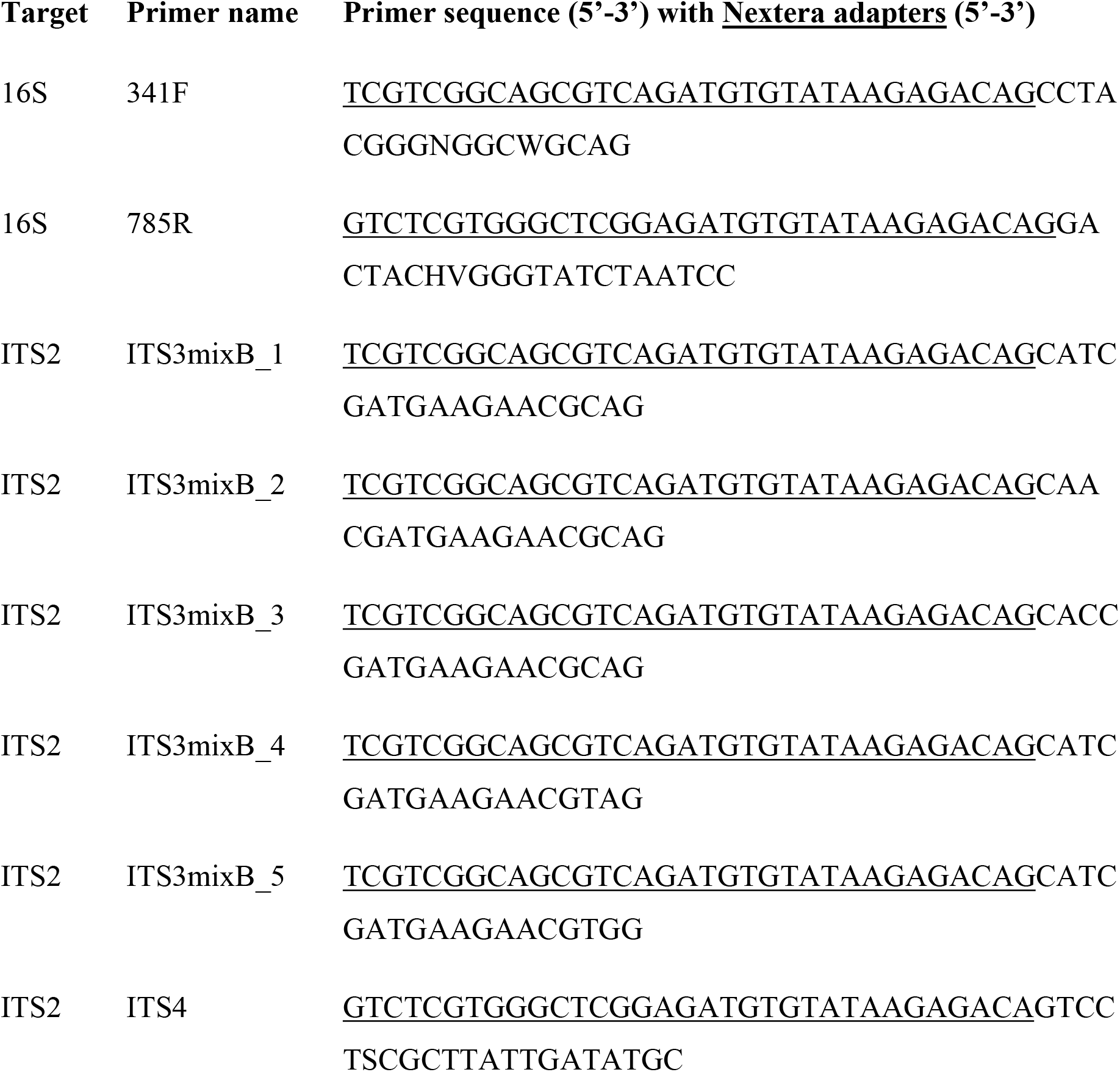
16S rRNA gene (v3-v4 hypervariable region) and ITS2 genomic region primers.

**Table S2.**
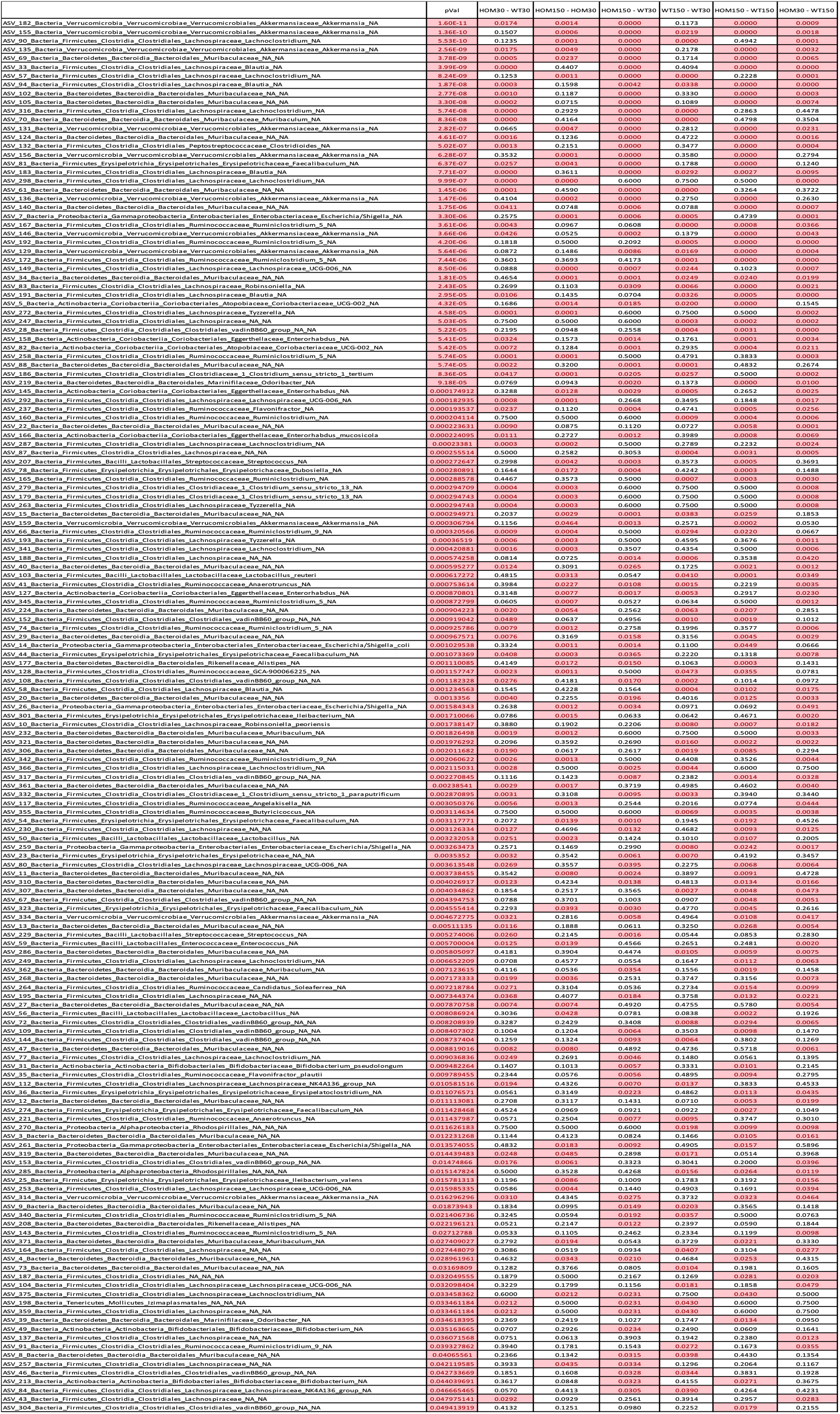
Bacterial ASV level analyses: Kruskal-Wallis rank sum test p-values represent difference between treatment groups and the Dunn test show multiple pairwise comparisons for each treatment group combination with fdr adjusted p-values. Values in red represent significant p-values (<0.05).

**Table S3.**
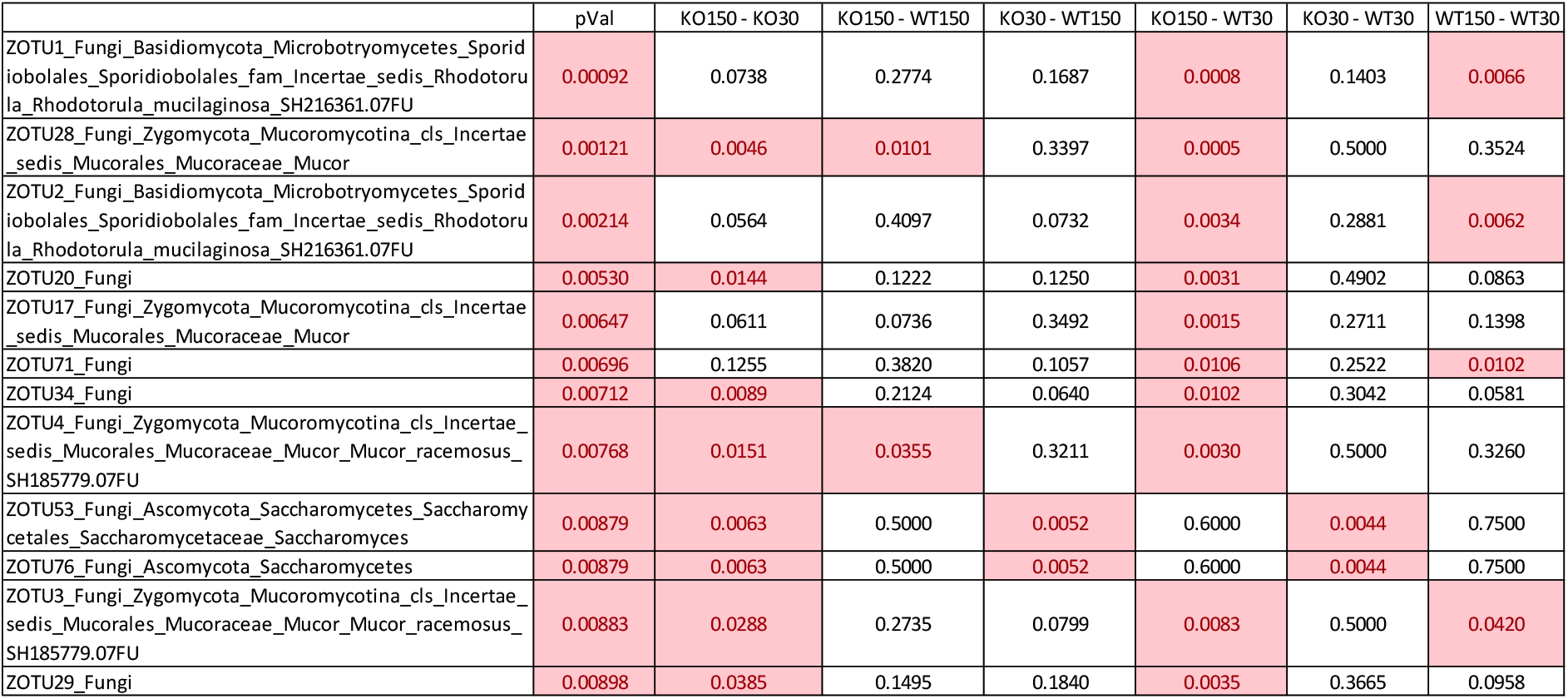
Fungal ZOTU level analyses: Kruskal-Wallis rank sum test p-values represent difference between treatment groups and the Dunn test fdr adjusted p-values from multiple pairwise comparisons for each treatment group combination. Values in red represent significant p values (<0.05).

**Figure S1.**
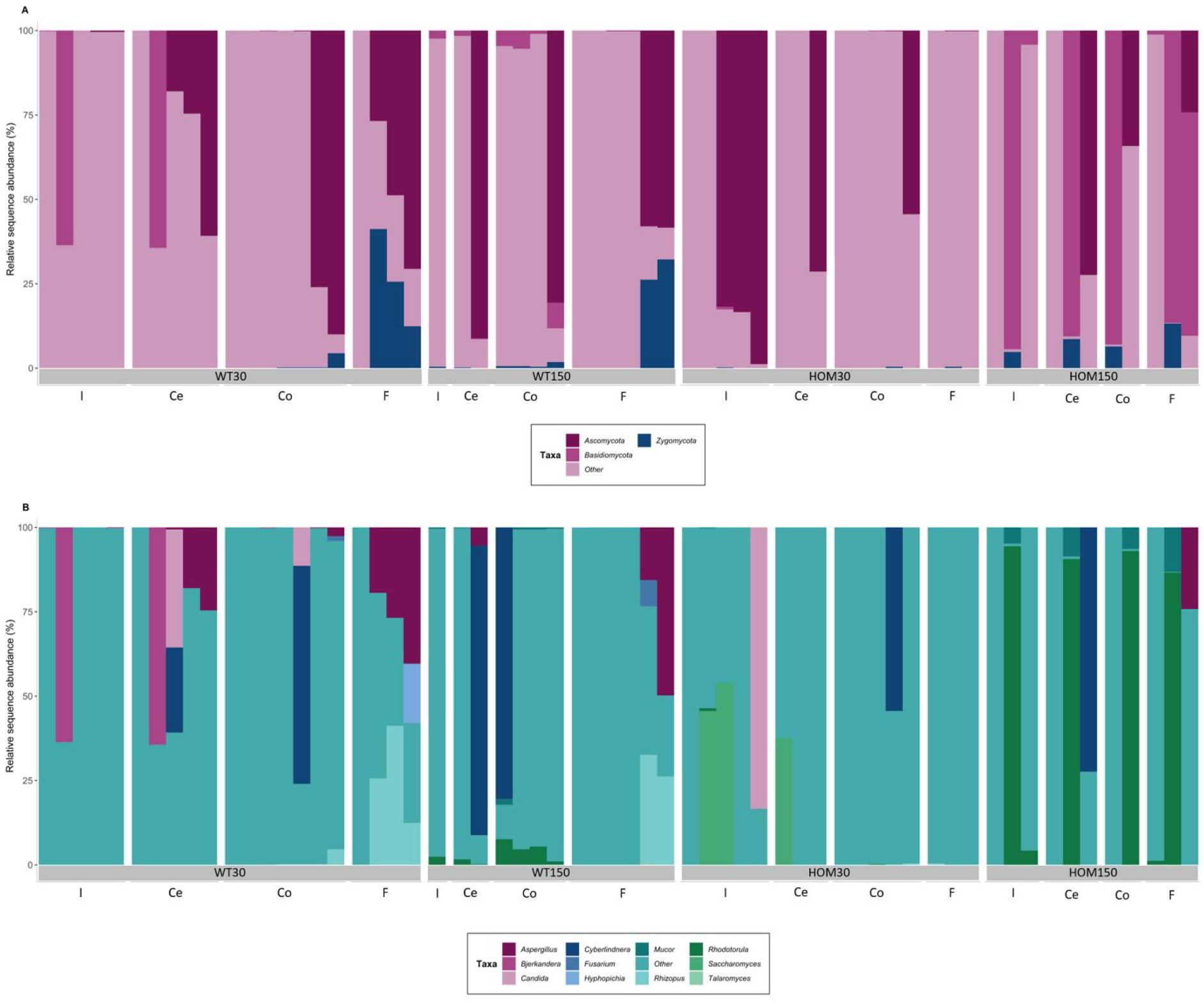
ITS2 taxonomic summary plot of the relative abundance of phyla (A) and top 20 most abundant genera (B) in each of the four treatment groups. (Wild type control zinc diet WT30 n = 21, wild type supplementary zinc diet WT150 n = 13, Shank3B−/− KO control zinc diet KO30 n = 16 and Shank3B−/− KO supplementary zinc diet KO150 n = 11). Gastrointestinal sections are noted as I (ileum), Ce (caecum), Co (colon), F (faecal).

